# Single cell RNA-seq reveals cellular diversity and developmental characteristics of human infant retina

**DOI:** 10.1101/617555

**Authors:** Jixing Zhong, Dongsheng Chen, Fangyuan Hu, Fang Chen, Zaoxu Xu, Jian Fang, Jianjiang Xu, Shiyou Wang, Langchao Liang, Chaochao Chai, Xiangning Ding, Jiacheng Zhu, Peiwen Ding, Jun Xia, Sanjie Jiang, Wei Li, Ya Gao, Jianguo Zhang, Jiankang Li, Ping Xu, Fengjuan Gao, Dandan Wang, Daowei Zhang, Jie Huang, Shoufang Qu, Shenghai Zhang, Jihong Wu, Xinghuai Sun

## Abstract

Retina, located in the innermost layer of the eye of human, holds the decisive role in visual perception. Dissecting the heterogeneity of retina is essential for understanding the mechanism of vision formation and even the development of central nervous system (CNS). Here, we performed single cell RNA-seq, analyzed 57,832 cells from human infant donors, resulting in 20 distinct clusters representing major cell types in retina: rod photoreceptors, cone photoreceptors, bipolar cells, horizontal cells, amacrine cells, Muller glia cells and microglia. We next constructed extensive networks of intercellular communication and identified ligand-receptor interactions playing crucial roles in regulating neural cell development and immune homeostasis in retina. Though re-clustering, we identified known subtypes in cone PRs and additional unreported subpopulations and corresponding markers in rod PRs as well as bipolar cells. Additionally, we linked inherited retinal disease to certain cell subtypes or subpopulations through enrichment analysis. Intriguingly, we found that status and functions of photoreceptors changed drastically between early and late retina. Overall, our study offers the first retinal cell atlas in human infants, dissecting the heterogeneity of retina and identifying the key molecules in the developmental process, which provides an important resource that will pave the way for retina development mechanism research and regenerative medicine concerning retinal biology.

## Introduction

Retina is the system responsible for light perception and visual signal transmission. Retina is a part of the central nervous system (CNS) and possesses diversified cell types. Thus, decoding the heterogeneity of retina is crucial for understanding the mechanism of vision formation as well as the highly complicated CNS. In general, seven major cell types (rod, cone, Müller glia cell, amacrine cell, horizontal cell, bipolar and retinal ganglion cells) constitute the three layers in retina: photoreceptors, bipolar cell layer and ganglion cell layer from inside out^1^. Photoreceptors, including rods and cones, are responsible for photo capture and convert them into electrical signal which will pass on to interneurons in bipolar cell layer for further modulation. Ganglion cells relay signals from interneurons to certain brain regions through optic nerves. Finally, the brain deciphers the electrical signals into three-dimensional vision. As the only non-neuronal cell, Muller glia cell is involved in extracellular environment regulation via nutrient intake, debris clearance, structural and functional stability maintenance^2,3^.

In recent years, the boost of scRNA-seq techniques allow us to zoom in to retina and have discovered tremendous subtypes in traditionally defined classification. In human, 20009 single cells from three post-mortem donors were divided into 18 distinct clusters, including well-known retinal cells and novel rod cells subpopulations^4^. In mouse, 40 subtypes of retinal ganglion cells and their markers were identified using scRNA-seq^5^. Developmental researches also shed lights on retina and CNS morphogenesis. A recent study in mouse analyzed cells from ten developmental stages and found that RPCs displayed a transcriptomic temporal transition between early stage and late stage. Also, NFI family of TF were designated as regulator of bipolar cells and Muller cells cell fate determination due to the enriched expression in late RPCs^6^. Another research characterized 24979 cells from human fetal retina from gestational weeks 12 to 27, revealing the differentiation programs of retinal cells and specifying the potential of early born Muller cells to be RPCs^7^.

Nevertheless, the retina development of postnatal infants, which impacts visual system vitally, remains elusive. In this research, we analyzed 57,832 single-nuclei transcriptomic profiles from infants in different development stages. Unsupervised clustering has defined 20 clusters, all of which were assigned with 7 major retinal cell types correspondingly. Communication network of 20 clusters connects the receptor-ligand pair with certain biological process in retina. We next performed reclustering thus providing a more delicate map of PRs and BCs. Reported subtypes of cone PR were confirmed and new subtypes of BCs were identified.

## Results

### Sample preparation and single cell transcriptome

We collected retinal biopsies from 2 donors (10-month-old and 2-year-old, hereafter termed as TenM and TwoY respectively) and dissociated them into single-cell or single nuclei suspension without surface marker pre-selection (Figure S1a). We constructed 3 snRNA-seq libraries for TenM retina and 2 scRNA-seq libraries for TwoY retina and then obtained the transcriptome of 66,362 nuclei/cells. After filtering, a total of 57,832 nuclei/cells were retained from subsequent analysis. The average number of genes detected per sample was 18,154, with a mean coverage of 57,545 reads per cell (Figure S1b). Pairwise Pearson correlation analysis demonstrated that libraries from the same donor share stronger correlation, confirming the validity of our datasets (Figure S1c). Of note, TwoYZJ and TwoY8 are standard scRNA-seq libraries thus explaining the relatively low correlations of these two libraries against others.

### Cellular heterogeneity in retina

We embarked on creating a map for human infant retina. In a global view, 57,832 cells were clustered into 20 clusters (Figure 1b). Batch effects from five different libraries were minimal (Figure S2a). According to canonical markers, differentially expressed genes (DEGs) and functional enrichment analysis (Figure 2c, f, g), we were able to identify several major retinal cell types and 1 microglia cells: Rod photoreceptors (C0, C1, C2, C3, C4, C5, C6), cone photoreceptors (C11, C17), bipolar cells (C8, C10, C15, C19), horizontal cells (C13), amacrine cells (C16), Muller glia (C7, C12, C18), microglia (C9, C14). Rod photoreceptors highly expressed *NR2E3* and *PED6A* and cone photoreceptors were characterized by the expression of *ARR3* and *CNGB3.* Bipolar cells exhibited specific expression of *TRPM1* while *GRIA3* for horizontal cells. Apart from the common glia cells marker (*SPP1*), Muller glia cells and microglia cells can be annotated clearly by the expression of *SLC1A3* and *HLA-DPA1* respectively. Additionally, functional related GO terms were enriched in corresponding clusters, further confirming the annotation result of each cluster. For example, photoreceptors clusters were strongly associated with sensory perception of light stimulus and visual perception while BC were enriched in GO terms related to modulation of chemical synaptic transmission. Microglia, known to be the primary resident immune cell type, displayed enrichment in neutrophil activation involved in immune response and T cell activation (Figure 2g). For ACs, though modest expression of corresponding markers were detected, hierarchical clustering and functional enrichment analysis exhibited AC characteristics in C16.

**Figure 1.**
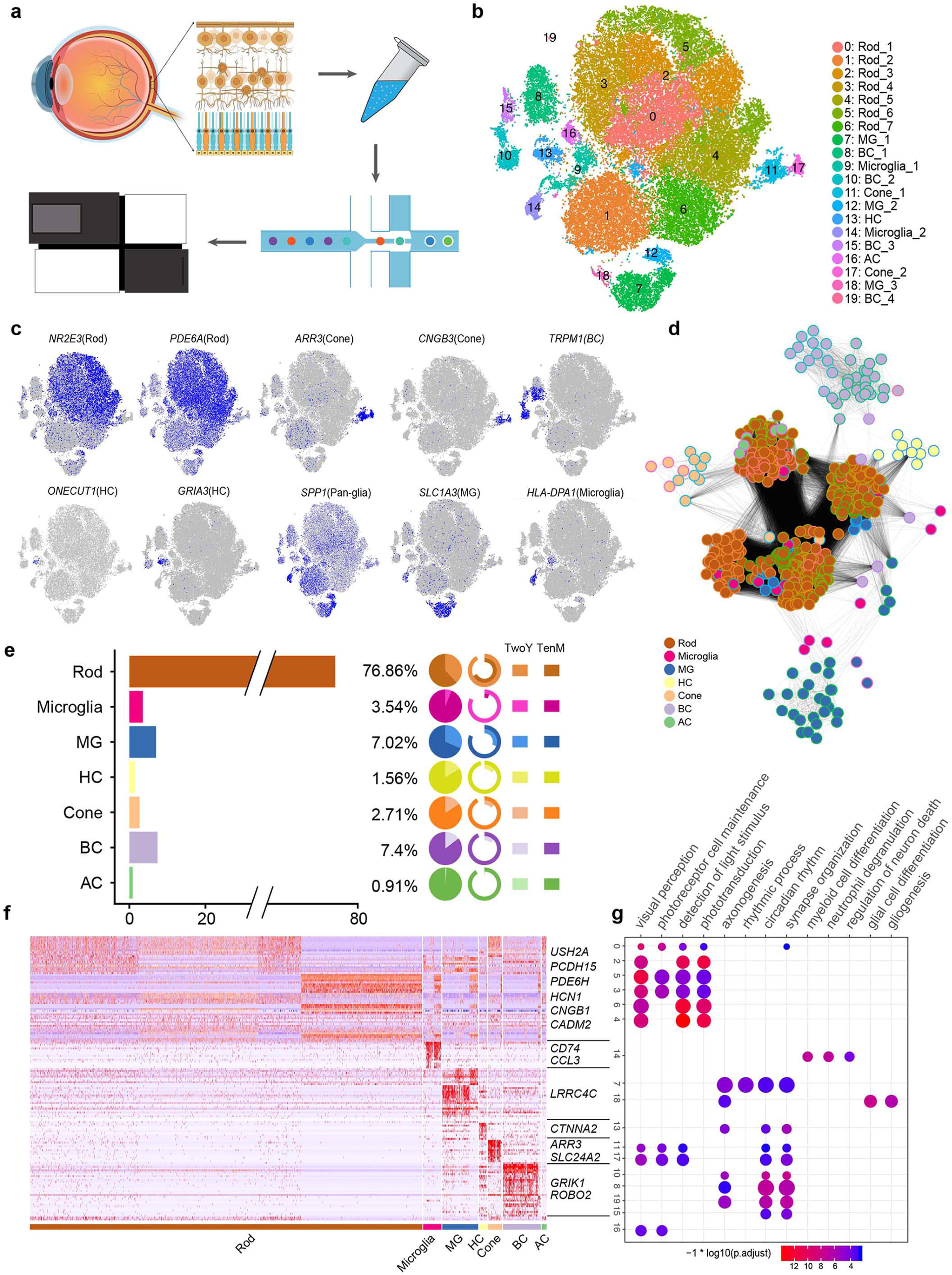
Cellular diversity in retina. (A) Scheme of the experimental and bioinformatics design in our study. (B) A t-SNE map of all libraries. Cells are colored by cell-type cluster. (C) T-SNE maps of all tissues single-cell data with cell colored based on the expression of marker genes for particular cell types. Gene expression levels are indicated by shades of blue. (D) A correlation network showing relationships among 20 cell-type clusters defined in Figure 2b. Each node represents the averaged expression of every 100 cells within each of the clusters. Each edge corresponds to a network correlation between two nodes. Each node is colored by corresponding cell type and node border is colored by cluster in Figure 2b. (E) Barplot showing the percentage of each cell type in the whole dataset. Pie charts: the proportion of cells from a certain sample in each cell type to the total number of cells identified as this cell types. Radial barplot: the proportion of cells from a certain sample in this cell type to the total number of cells in this sample. (F) A gene expression heatmap showing top differentially expressed gene for each cluster in all 20 clusters. Red corresponds to high expression level; white indicates intermediate level and blue correspond to low expression level. (G) Selected GO terms enriched in each cellular cluster. Dot size denotes the gene ratio and color denotes the adjusted P value.

**Figure 2.**
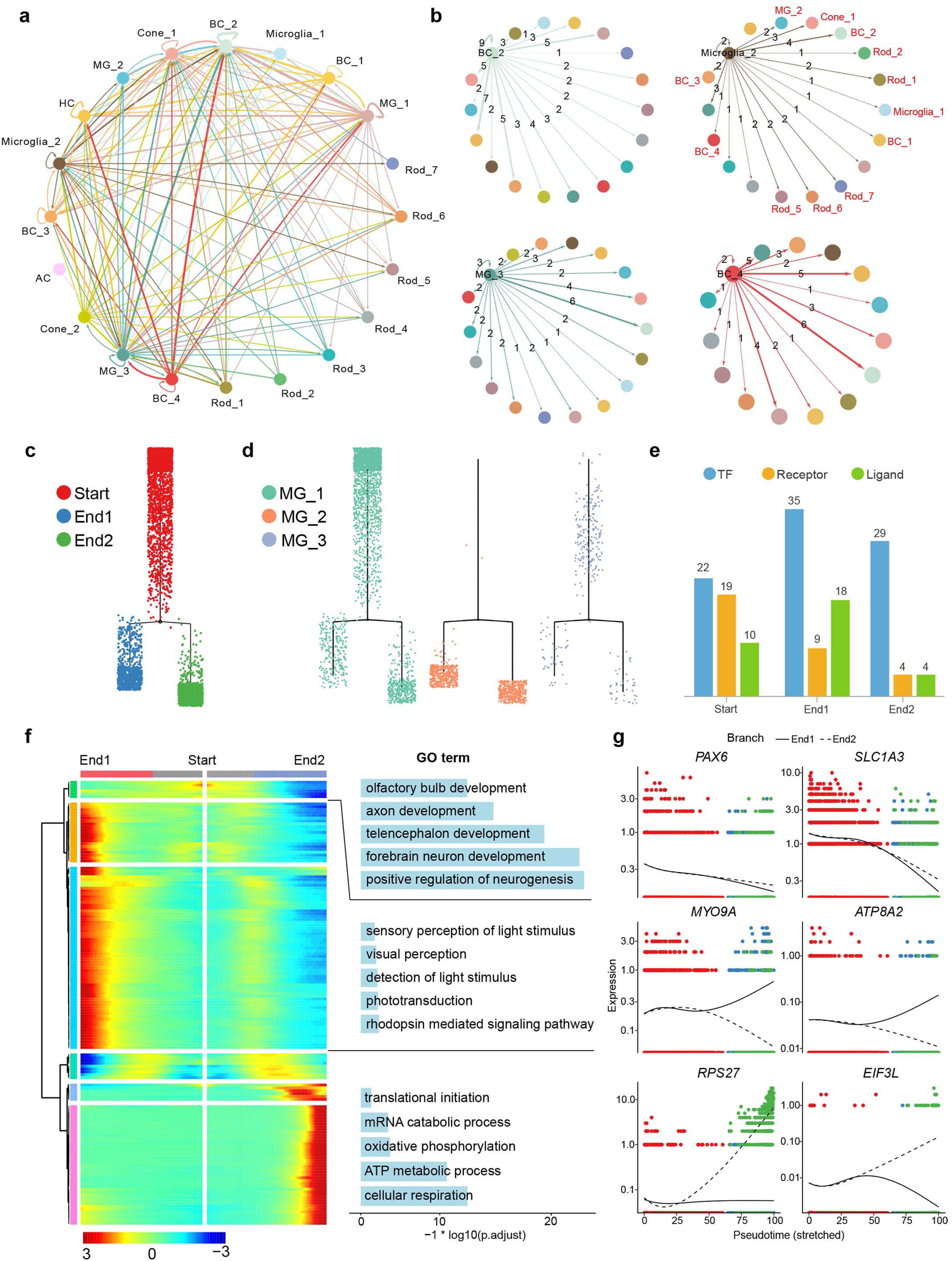
(A) Intercellular communication within each cluster in Figure 2b. The line color indicates ligands broadcast by the cell population of the same color (labeled). Lines connect to cell populations where cognate receptors are expressed. The line thickness is proportional to the number of ligands where cognate receptors are present in the recipient cell population. Lines with number of interactions lower than 1 are not shown. Map quantifies potential communication, but does not account for anatomic position or boundaries of cell populations. (B) Sub-communication network of selected clusters. (C-D) Cell lineage relationships of Muller cells (MG_1, MG_2 and MG_3 in Figure 1b) in the human infant retina revealed by Monocle branched single-cell trajectory analysis. Each dot represents a single cell; the color of each dot indicates the corresponding branch information assigned by Monocle(C) or the cell type information (D). (E) Barplot showing the numbers of differentially expressed TF, receptor and ligand in each branch state. (F) Gene ontology analysis of modules created by clustering the two main branches from the lineage tree. The analysis reflects the cell fate commitment. In this heatmap, the middle represents the start of pseudo-time. One lineage starts from the start and moves to the End1 while the other lineage moves to the End2. Rows are genes correlated into different modules. (G) Expression of selected differentially expressed genes for each branch. The expression of cells was ordered by Monocle analysis in pseudo-time. The full line represents the End1 while the dash line represents the End2.

To systematically dissect the relationships between different cell types, we built correlation-based networks at the cell-type level (Figure 1d). We binned cellular cluster into sub-groups with 100 cells to reduce noise and enhance the robustness of the network. Consequently, the correlation map mimicked a cellular landscape for different retinal cell types. In general, the majority of cells showed much denser sub-network with cells from the same cell type than with others. Notably, almost all cells showed connections with rod PR, revealing high correlation with rod PR at averaged transcriptomic level. Another interesting observation was that, glia cells, including microglia cells and part of Muller cells, dispersed amidst rod PRs instead of forming distinct sub-network.

In order to assess the cellular composition in retina, we calculated the percentage of each cluster in the whole dataset (Figure 1c). The constitution of infant retina showed high similarity when compared with human adult retina dataset^4^. Rod photoreceptors took up the majority (75.32%) of the whole dataset while cone photoreceptors occupied a very small portion (3.1%). The second-largest cell type is Muller glia (9.24%). However, amacrine cells which constituted ∼10% of the total cells in mouse, only account for 0.92% of our data. We also investigated the proportion of each cell type in early and late retina sample, which are shown in pie chart (absolute number) and radial barplot (proportion) (Figure 1c).

We next moved on to explore the intercellular communication network within retina. The cellular interactions were inferred using published ligand-receptor pairs (Methods). Expression patterns of ligand-receptor pairs in the networks revealed dense intercellular communication networks, especially in glia and BCs (Figure 2a). Nevertheless, rod PR subpopulations exhibited a relatively sparse network, showing less than one interaction among subpopulations within themselves and few with other cell types. This indicated that rod PRs interacted much more often with other retinal cell types than with PRs subpopulations. The most frequent interaction that appeared in our dataset is RTN4— LINGO1 which is suggested to regulate the radial migration of cortical neurons in mouse brain^8^, implying that RTN4—LINGO1 may be involved with migration of neural cells in retina. NRG1-ERBB4 and NRG3—ERBB4 pairs were also extensively expressed in retina neural cells—Muller glia interactions. *NRG3* has been reported to play a key role in regulating the growth and differentiation of glia cells and its signal transduction with ERBB4 was essential for neuron circuit formation^9^ (Figure 2b). *SEMA3A*, specifically expressed in rod PR (C0, C16) can signal through PLXNA4 and engage in multiple biological activities like axon pruning and dendrites branching in adult procine^10^ (Figure 2b). Particularly, Microglia_1 (C14) expressed HLA-A and its cognate receptor APLP2 were detected in the majority clusters (C0, C1, C4, C5, C6, C8, C9, C10, C11, C12, C15, C19), covering PRs, BCs and Muller cells (Figure 2b).

### Developmental trajectory of Muller glia

Muller glia, as the most common type of glia cell in retina, serve to maintain the structural and functional stability of retinal cells. Previous study has reported that Muller glia showed similar properties as retinal progenitor cells^7^ and were capable to differentiate into photoreceptors in human and mouse cell culture^3,11,12^.

To further investigate the developmental relationship of MG clusters identified in Figure 1b, we subset MG and performed pseudo-time trajectory reconstruction using Monocle. Consequently, we obtained a putative developmental path with two branches (Figure 2c). We then labeled MG subpopulations in the trajectory and found that MG_1 were primarily centralized on the start, MG_2 converged on both ends while MG_3 distributed loosely along the developmental path (Figure 2d). In order to explore the different properties of cells located in the starting point and two branches of the trajectory, we adopted branch expression analysis modeling (BEAM, Methods) to calculate DEGs. GO term enrichment analysis revealed that genes upregulated in cells in the start of the path is associated with neurogenesis and neuron development (Figure 2f). *PAX6*, a key regulator in the development of retinal neural tissues, showed relatively high expression in the start of the path and declined along with the path proceeding (Figure 2g). Cells in End1 were enriched with GO terms related to light perception whereas cells in End2 displayed features of active translation and respiration processes (Figure 2f). On top of that, cells in End1 showed *ATP8A2* gene expression, which was reported to be required for normal visual function and involved in photoreceptor survival^13^. *RPS27*, which encodes a ribosomal subunit protein and *EIF3L*, which specifically targets and initiates translation of a subset of mRNAs involved in cell proliferation and cell cycling^14^, were highly expressed in cell in End2 (Figure 2g). These results implied the distinct states of cells in two Ends.

### Heterogeneity within major cell types: PRs and BCs

Due to the high heterogeneity of retinal cells, we next provided a more specific map for major cell types in retina by reclustering of corresponding clusters.

Based on previous study, there are three types of cone photoreceptors in human retina that can be differentiated by the expression of three opsins (OPN1SW, OPN1MW, OPN1LW), responsible for the perception of colors with different wavelengths^15,16^. To assess the heterogeneity of cone, we performed reclustering in cone PR clusters and divided them into 3 subclusters (Figure 3a). Due to the highly homologous sequence between *OPN1MW* and *OPN1LW*, we were only capable of distinguishing the S-cones (C2) from M/L-cones (C0, C1) using opsins encoding genes (Figure 3c). In this cone PR dataset (1569 cone PRs), M/L-cones constitute 94.77% of the total cells and the remaining 5.23% are S-cones (Figure 3b), which was a bit higher than the proportion (3.19%) from a scRNA-seq study into human adult retina^4^. Next, we conducted gene differential expression analysis to reveals new genetic characteristics for S-cones and M/L-cones. Highly and specifically expressed genes for S-cones (e.g. *KIAA1217, VWC2, SMAD5*) and M/L-cones (e.g. *RPS16, DDX5, SMTN*) were identified (Figure 3d, Table S2), providing more molecular evidences for future cone PRs studies.

**Figure 3.**
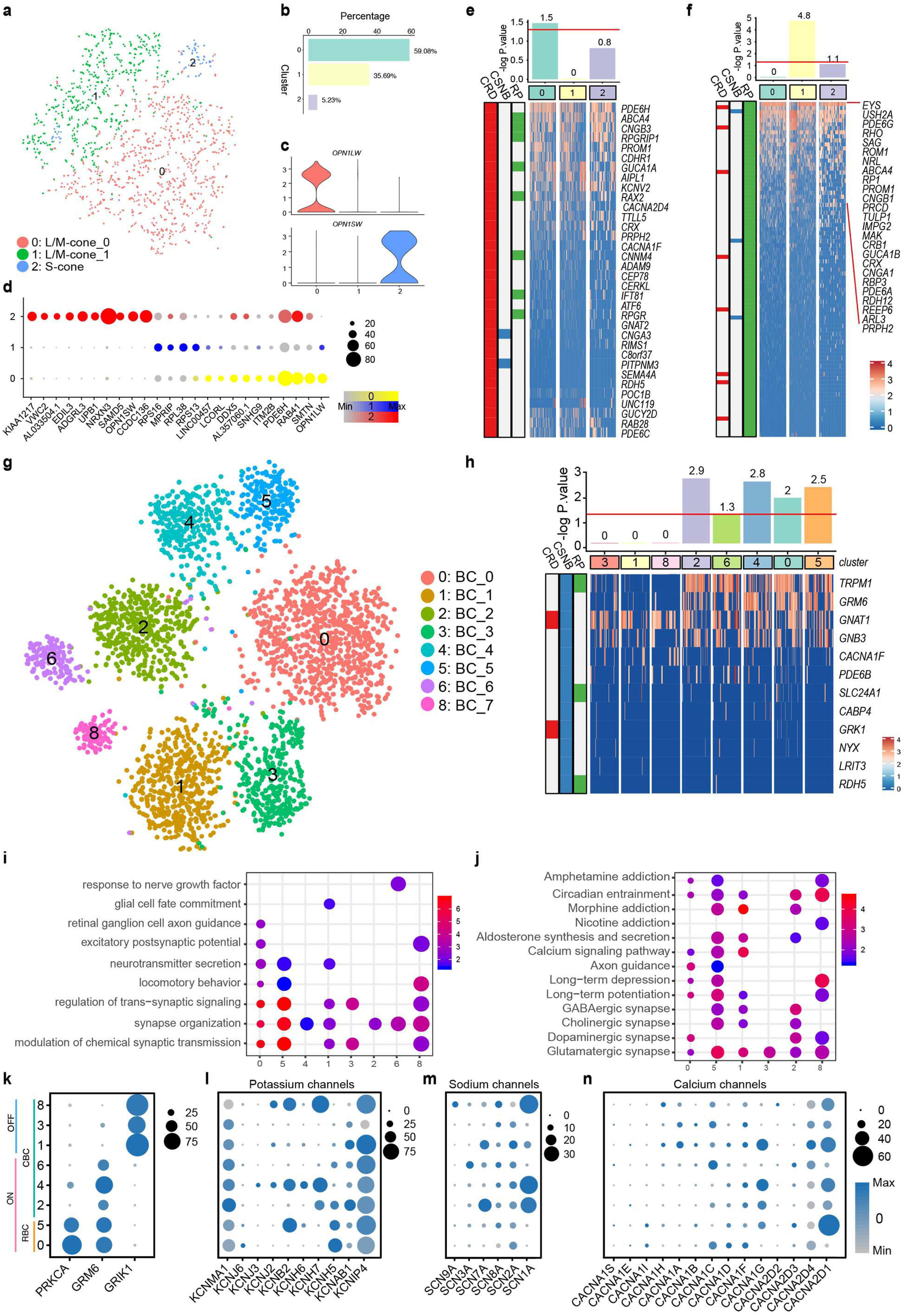
Cellular diversity within cone PRs and BCs. (A) A t-SNE map of cone photoreceptors single-cell data. Cells are colored by cluster (subtypes). (B) Barplot showing the percentage of S-cones and M/L-cones in the whole cone PR dataset. (C) Violin plot displaying the expression of opsin encoding genes in each cluster in Figure 3a. (D) Bubble plot showing the expression of differentially expressed genes in each cluster in Figure 3a. Dot size represent the percentage of cells express corresponding gene and color represent the expression level. (E,F,H) Cone subtype enrichment of CRD (Cone-Rod Dystrophy) (E), RP (Retinitis Pigmentosa) (F) and bipolar subtype enrichment of CSNB (Congenital Stationary Night Blindness) (H) risk genes respectively. Heatmap: Expression of corresponding disease risk genes in cell subtypes. cells are ordered by cluster. Bar graphs: numbers indicate log2 P value, the red line indicates significance. (G) A t-SNE map of bipolar cells single-cell data. Cells are colored by cluster (subtypes). (I) Enriched GO terms in each cluster in Figure 3f. Dot size denotes the gene ratio and color denotes the adjusted P value. Dot size represent the percentage of cells express corresponding gene and color represent the expression level. (J) Enriched KEGG pathways in each cluster in Figure 3f. Dot size denotes the gene ratio and color denotes the adjusted P value. (K) Expression pattern of BC subtypes classification genes. (L-N) Bubble plot showing cluster-specific expression of Potassium (L), Sodium (M) and Calcium (N) channels encoding genes.

We next interrogated another kind of PR in retina — rod PRs. On average, there are approximately 92 million rod cells in the human retina^17^. Rod cells are more sensitive than cone cells and are almost entirely responsible for night vision. In our dataset, 44448 rod cells were divided into 7 subpopulations (Figure 1b, Figure S7a). To investigate the sample origin of each cluster, we calculated the proportion of cell derived from the early and late retina in each cluster (Figure S7b). We found that Rod_1 and Rod_2 subpopulations were dominant by cells from early retina while the rest subpopulations were primarily comprised of cells from late retina. To assess the similarity of clusters at transcriptomic levels, we performed hierarchal clustering based on transcriptomic data. DEGs expression patterns and corresponding GO term enrichment results suggested the different features of rod subpopulations (Figure S7c,d). Bipolar cells, served as interneurons, receive electrical impulses from photoreceptors, process them in various manners and then pass them to RGC. The diverse subtypes of BCs ensure their multi-functioning roles. Previous researches confirmed 15 subtypes of BC experimentally using ∼25,000 BCs^18^. BCs can be divided into two categories according to the type of photoreceptor which they connect to. Another taxonomy classifies BCs into ON and OFF types based on whether they will be excited or suppressed by increasing illumination. In our BC dataset (4282 BCs), unsupervised reclustering produced 9 distinct subclusters (Figure 3g). Of note, C7 were thought to be photoreceptor contamination and excluded from downstream analysis due to the modest expression of BC markers and light perception concerning GO terms enriched using DEGs of C7 (Figure S6b, c). C7 also displayed higher correlation with rod subtypes instead of BC subtypes (Figure S6d). When we took a closer look into C7 we found that cells in C7 were scattered in the whole BC clusters in Figure 2a (i.e. C8, C10, C15, C19 in Figure 2a). One possible explanation is that reclustering of BC dataset puts photoreceptors into a BC environment, extending the differences when it compared to the differences in the whole dataset situation. C0 and C5 were annotated as rod bipolar cells (RBC), showing high expression of *PRKCA*. According to the clear expression pattern of *GRM6* and *GRIK1*, C0, C2, C4, C5, C6 were considered as ON BCs while the remaining C1, C3, C8 were OFF BCs (Figure 3k). Heatmap of DEGs displayed distinct expression pattern of the defined clusters. We further explore the function of DEGs by functional enrichment analysis (Figure 3i, j). GO terms that closely related to the signal transduction like modulation of chemical synaptic transmission, neuron recognition and morphogenesis like axon guidance and synapse organization were enriched in almost all clusters to a varying extent. Next, we sought to fully describe the type-specific functional roles of BC subtypes by assessing the expression of Sodium, Calcium and Potassium ion channel subunits because ion channels control the transmit of neural signals. Sodium channel subunit-encoding genes were expressed selectively among subtypes (Figure 3l-n). Expect for *SCN1A*, *SCN2A, SCN3A* and *SCN8A* that were previously reported to be expressed in CBCs in goldfish^19^, we noticed that *SCN7A* and *SCN9A* also showed specific expression among CBCs. Calcium channel alpha and delta subunits encoding genes also showed varying expression levels among subtypes. Furthermore, we observed a great deal of differentially expressed TFs, receptors and ligands, supporting our deductive functional subtypes (Figure S6e).

### Cell type enrichment analysis of retinal diseases associated genes

We retrieved retinal diseases causing genes from RetNet database (https://sph.uth.edu/retnet/disease.htm) (Table S4) and performed hypergeometric test to see whether these disease-related genes are enriched in the DEGs of certain clusters (Methods), by which means, discovering putative disease-causing cell type and therefore providing new sights for retinal diseases. Most of the disease types we investigated in this study are caused by malfunctions of PRs. Retinitis pigmentosa (RP) is characterized by retinal cells degeneration, begun with rod cells and followed by cone cells, consistent with which, enrichments of RP related genes occur in all rod subpopulations and M/L-cones (Figure 3f, Figure S7f). This indicated that all rod subpopulations but only M/L-cones are involved with RP genesis and corresponding markers we identified may help to pinpoint the scope of target cells for therapies. Cone-rod dystrophy (CRD), unlike RP, is featured by an initial loss of color vision and of visual acuity due to loss of cone function and followed by rod cells deterioration. Most rod subpopulations (Rod_1, Rod_2, Rod_5, Rod_6, Rod_7) and M/L-cones displayed close connections with CRD occurrence with enrichment significance (-log P value) over 1.5 (Figure 3e, Figure S7e). Congenital stationary night blindness (CSNB) can be classified into complete CSNB (cCSNB) and incomplete CSNB (iCSNB) which is caused by defects localize to ON BCs and PR synapse respectively. Accordant with what mentioned above, we found that ON BCs DEGs are highly enriched in CSNB causing genes, but to a substantially varying extent (Figure 3h), suggesting that different BC subtypes contribute differently to CSNB or lead to CNSB with various severities. Furthermore, DEGs of Rod_1, Rod_2, Rod_5 and Rod_7 subpopulations showed modest association with CSNB causing genes (Figure S7f), implying a moderate role in CNSB. Besides, macular degeneration (MD) were specifically associated with Rod_1, Rod_2 and Rod_7 (Figure S7h). Together with what described above, Rod_1, Rod_2 and Rod_7 showed strong enrichment of common retinal diseases. From the perspective of cone PRs, S-cones don’t seem to be significantly related to any retinal disease we collected while M/L-cones are the major cone PR subtype causing or being affected by retinal diseases.

In conclusion, these results not only matched previous findings but also linked several diseases with certain retinal cell subtypes, facilitating researches and development of novel drug into retinal malfunctions.

### Similarities and differences in cell proportion, cell status between retina in different developmental stages

Since cell proliferation is a key metric of cell maturation level, we next performed cell cycle analysis. We first assigned a cell cycle phase score to each cell using gene expression signatures for the S and G2/M phases. From a global view, most of the cells were in G1 phase while relatively few were in proliferating G2/M stage (Figure S9a). Interestingly, cells in G2M stage were primarily from early retina (Figure S9c). Except for rod PR where cells from late retina diffuse more evenly, most cells from late retina in other cell types were assigned negative S scores and G2M scores, thus being considered to be in G1 phase.

We next asked what programs the postnatal retina will go through. Previous studies have shown that photoreceptors will develop from a relatively immature state into a mature state, improving the light sensitivity after birth^20^. Hence, to answer the question above, we performed differential expression analysis in the same cell type between early and late PRs.

In cone PRs, 581 upregulated and 920 down-regulated genes in total were found in early cone RPs in M/L-cones (C0, C1 in Figure 3a, Table S2) when compared with late cone RPs (Figure 4a, c). Functional enrichment demonstrated that genes upregulated in early cone PRs showed weak enrichment in terms related to light perception while highly express genes concerning photoreceptor differentiation (Figure 4b, d). In S-cones (C2 in Figure 3a), 44 down-regulated whereas no upregulated genes found to be significant (Figure S8h, Table S2). GO terms associated with cellular respiration were enrichment in genes upregulated in late PRs (Figure S8h). We next inspected those TFs, receptors and ligands that differentially express between two stages in M/L-cones (Figure 4e). We found that *ARID1B*, which were highly expressed in early cone PRs, belongs to a neuron-specific chromatin remodeling complex and involved in cell-cycle activation and neural progenitor cell fate commitment during neural development^21^. *MEF2C*, also displayed specific expression in early cone PRs, has been prove crucial for normal neuronal development, distribution, and electrical activity in the neocortex^22^. *NR2E3*, an orphan nuclear receptor of retinal photoreceptor cells, is an activator of rod development and repressor of cone development^23^. Hence, it’s reasonable to infer that TFs like *ARID1B, MEF2C* and *NR2E3* cooperate together to drive the development process in cone maturation. Likewise, in rod PRs, according to the functional enrichment analysis, it’s also showed that cells from early retina displayed strong differentiation ability while cells from late retina exhibit the competence to perceive external light and transduce signal to subsequent neurons (Figure S8).

**Figure 4.**
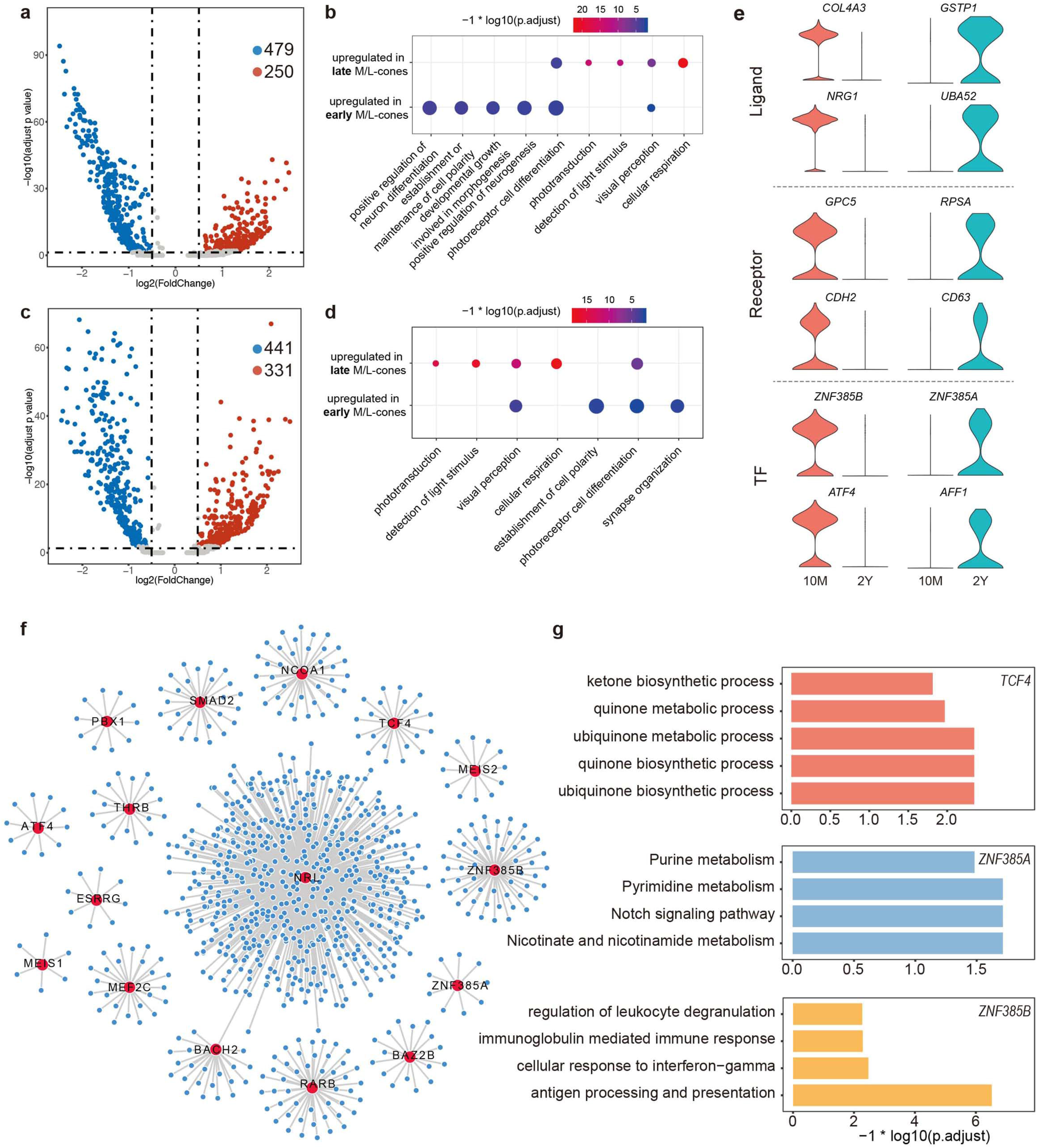
Developmental analysis of cone PR from early and late retinas. (A) Volcano plot showing the differentially expressed genes in M/L-cone_0 (Figure 3a) between early and late retina. Genes are plotted as log2 fold change versus the −log10 of the adjusted p-value. Genes in red denotes genes upregulated in early retina while blue denotes upregulated in late retina. Insignificant genes are colored in grey. Thresholds are shown as dashed lines. (B) GO term enrichment analysis of differentially expressed genes in M/L-cone_0 between early and late retina. (C) Volcano plot showing the differentially expressed genes in M/L-cone_1 (Figure 3a) between early and late retina. Genes are plotted as log2 fold change versus the −log10 of the adjusted p-value. Genes in red denotes genes upregulated in early retina while blue denotes upregulated in late retina. Insignificant genes are colored in grey. Thresholds are shown as dashed lines. (D) GO term enrichment analysis of differentially expressed genes in M/L-cone_1 between early and late retina. (E) Violin plot Differentially expressed TFs, receptors and ligands in M/L-cone_0 between early and late retina. (F) Regulatory network constructed using genes upregulated in M/L-cone_0 in early retina. Each node represents a gene and each line represents a putative interaction. Nodes colored in red represent TF and nodes colored in blue represent target genes. (G) GO term enriched in the target genes of TCF4, ZNF385A and ZNF385B.

To further identify key regulators in the course of cone PR development, we next constructed regulatory network based on significantly high expressed TFs in early M/L-cones_1 (C1 in Figure 3a) (Figure 4f). NRL, reported as a critical intrinsic regulator of photoreceptor development and function^24^, possessed most putative target genes, indicating its crucial role in the developmental course of cone PR. The targets of TCF4 were enriched in GO terms related ubiquinone and ketone biosynthetic processes (Figure 4g). Ubiquinone is the oxidized form of CoQ10, which serves as an antioxidant to help protect cells from oxidative damage. The level of CoQ10 declines with age due to decreased synthesis and increased degradation, leading to age-related neurodegenerative diseases, such as Parkinson’s disease and age-related macular degeneration^25^. Besides, ZNF385B, whose targets were found to be associated with immune activities, were reported to be expressed in germinal center B cells and involved in B-cell apoptosis^26^.

## Discussion

This complex intrinsic composition of retina cells highlights the necessity of single-cell technologies for dissecting transcriptome signature in detail. Here, we report the first single cell atlas – to our knowledge, for human retina in different infant developmental stages. Our dataset consists of 57,832 cell and nuclei achieve unprecedentedly large scale for human retina research. In this study, we firstly established a high-resolution map for human infant retina with six out of seven major retinal cell types successfully assigned. We also identified microglia, the resident immune cells in human retina and found by ligand-receptor communication network analysis that microglia showed putative interactions primarily with rod PRs, BCs and Muller glia.

Due to the progenitor-like features of Muller glia, we performed pseudo-time trajectory analysis using MG_1, MG_2 and MG_3, which revealed a developmental path with two branches. Further characterization of the branches of the path suggested that cells in left branch displayed light perception functions and cells in the right branch showed active translation activities.

We next sought to identified subtypes and subpopulations of photoreceptors and bipolar cells through reclustering. For cone PRs, we identified S-cones but failed to distinguished M-cones from L-cones due to the high homology of opsin sequences. For BCs, reclustering resulted in 8 subpopulations, 3 of which were classified as OFF BCs and the remaining 3 were thought to be ON BCs according to the expression of *GRIK1* and *GRM6*. We further described the different properties of each subpopulations by assessing the expression pattern of function related ion channels.

Another part of this study is to compare the similarities and differences of early and late retinas at transcriptomic level. We first evaluate the proliferation state of each cell by assigning each cell a calculated cell cycle stage. We found that except for microglia, most of the cells from late retina remain in a relatively quiescent in other cell types we identified. That is to say, cells from early retina were in G2M stage, an actively proliferating cell cycle stage. In addition, through differential gene expression analysis and functional enrichment analysis, we found M/L-cones from early retina showed higher expression of genes related to photoreceptor differentiation and neurogenesis whereas M/L-cone from late retina showed mature photoreceptor properties with enriched GO terms concerning light perception and phototransduction.

Inherited retinal diseases (IRDs) are a group of retinal disorders caused by an inherited gene mutation and can result in vision loss or blindness^27–29^. Currently, the disease diagnosis and therapies were hampered by our understanding of human retina at single cell level. In our retinal disease analysis, S-cones manifested insignificant enrichment of all common retinal diseases included in our analysis (Table S4) while M/L-cones were found to be associated with CRD and RP. For rod PRs, Rod_1, Rod_2 and Rod_7 showed strong association with many retinal genetic disease, especially with CRD and RP. CSNB were found to be associated with defects in ON BCs but the enrichment varies among the ON BC subpopulations. Taken together, our findings narrow down the disease-causing cell subtypes and subpopulations and their corresponding biomarkers, which may not only shed lights on understanding the molecular mechanisms underlying IRDs, but also provide new insights for discovering novel drug targets and developing gene therapies for IRDs.

## Material and methods

### Ethics statement

Collection of human retina was approved by the Ethics Committee of the Eye & ENT Hospital of Fudan University and in accordance with the Code of Ethics of the World Medical Association (Declaration of Helsinki) for medical research involving human subjects. All protocols were in compliance with the ‘Interim Measures for the Administration of Human Genetic Resources’ administered by the Chinese Ministry of Health.

### Human retina collection

The retinal tissues come from a 10-month and a 2-year-old toddler donor. The donor’s eyeballs were collected by Eye & ENT Hospital of Fudan University for corneal transplantation, and eyeball samples acquired within 12 hours post-mortem. The remaining eyeball tissue was used to extract retina tissue. Retina tissue could be exposed by removing iris, lens and vitreous successively. The intact retinal tissue was carefully dissociated and placed in pre-cooled HBSS without RPE/choroid layer. One of the retinal tissues was digested into single-cell suspension in Trypsin-EDTA solution (containing 0.25% Trypsin and 0.02%EDTA) for 37 °C 20 min, and the remaining tissues were immediately frozen in liquid nitrogen. Then, single-cell suspension was filtered through a 40μm cell strainer to eliminate any clumped cells. The freshly prepared cell suspension was immediately used for the construction of single-cell sequencing libraries.

### Single cell cDNA library preparation and high-throughput sequencing

The freshly prepared cell suspensions to generate cDNA libraries with Single Cell 3’ Library and Gel Bead Kit V2(120237 10x Genomics), according to the manufacturer’s instructions (120237 10x Genomics). Flash-frozen tissues were homogenized in 2mL ice-cold Lysis buffer (10 mM Tris-HCl, pH 7.4, 10 mM NaCl, 3 mM MgCl2, 0.1% NP40, protease inhibitors) then dounced in an RNase-free 2ml glass dounce (D8938-1SET SIGMA) 15x with a loose pestle and 15x with tight pestle on ice. Transfer homogenization through 40 μm filter (352340 BD) and removed the block mass. The cell filtrate was proceeded to Density Gradient Centrifugation. Mixed 400 μL cell filtrate with 400 μL 50% Iodixanol Solution in 2mL lo-Bind tubes (Z666556-250EA SIGMA). Carefully layer 29% Iodixanol Solution and 35% Iodixanol Solution to the bottom of tube. Centrifuging at 3,000 xg, 4°C for 30min.

Nuclei were resuspended in ice-cold 1 x PBS (10010-031 GIBCO) containing 0.04% BSA and spin down at 500xg for 5min. Discarded supernatant and then using regular-bore pipette tip gently pipette the cell pellet in 50 μL ice-cold 1 x PBS containing 0.04% BSA and 0.2U/μl RNase Inhibitor. Determined the nuclei concentration using a Hemocytometer (101010 QIUJING). Then, loaded on a Chromium Single Cell Controller (10x Genomics) to generate single-cell Gel Bead-In-EMulsions (GEMs) by using Single Cell 3’ Library and Gel Bead Kit V2 (120237 10x Genomics). Captured cells released RNA and barcoded in individual GEMs. Following manufacturer’s instructions (120237 10x Genomics) libraries were generated from each donor sample. Indexed libraries were converted by MGIEasy Lib Trans Kit (1000004155, MGI) then sequenced on the MGISEQ 2000 (MGI) platform with pair-end 26bp+100bp+8bp (PE26+100+8).

### Pre-processing and quality control of scRNA-seq data

We first used Cell Ranger 3.0.2 (10X Genomics) to process raw sequencing data and then Seurat (10.1038/nbt.3192) was applied for downstream analysis. Before we start downstream analysis, we focus on four filtering metrics to guarantee the reliability of our data. (1) We filter out genes that are detected in less than 0.1% of total cell number to guarantee the reliability of each gene; (2) We filter out cells whose percentage of expressed mitochondrial genes are greater than 10%; (3) We also filter out cells whose UMI counts are either less than 1.5 offset to the first quantile or greater than 1.5 offset to the third quantile of total UMI counts to filter out the doublet-like cells; (4) We filter out cells whose detected genes are less than 200.

### Identification of cell types and subtypes by dimensional reduction

The heterogeneity of retina was determined using Seurat R package (10.1038/nbt.3192). We performed Seurat-Alignment to eliminate batch effect, allowing us to combine data from multiple samples. Then we determined significant PCs using the JackStrawPlot function. The top twelve PCs were used for cluster identification with resolution 1.0 using k-Nearest Neighbor (KNN) algorithm and visualization using t-Distributed Stochastic Neighbor Embedding (tSNE) algorithm. Cell type were assigned by the expression of known cell-type markers and functional enrichment analysis. The FindAllMarkers function in Seurat was used to identify marker genes for each cluster using default parameters.

### Removal of cell cycle effects in clustering and cell cycle analysis

We collected 43 genes and 54 genes related to S phase and G2/M phase respectively^30,31^. For clustering, each cell was assigned a score to delineate its cell cycle state by CellCycleScoring function in Seurat according to the expression of these genes. Subsequently, the cell cycle effect was regressed out based on the scores, leading to a more accurate clustering result. For cell cycle analysis, cells were determined to be quiescent (G1 stage) if their S score < 0 and G2/M score < 0; otherwise, they were deemed proliferative. In addition, proliferative cells were designated G2/M if their G2/M score > S score, whereas cells were designated S if their S score > G2/M score.

### Functional enrichment analysis

Functional enrichment analysis includes two parts: GO term and KEGG pathway. Lists of genes were analyzed using clusterProfiler R package (10.1089/omi.2011.0118) and the BH method was used for multiple test correction. GO terms with a P value less than 0.01 and KEGG term with a P value less than 0.05 were considered as significantly enriched.

### Construction of intercellular correlation network

To reduce noise, we averaged the expression of every 100 cells within cluster and then calculated the pairwise Pearson correlation between two dots based on their average expression profiles. Inter-dots relationship will be shown if their Pearson correlation is greater than 0.8. The visualization of the correlation network is achieved using Cytoscape (10.1101/gr.1239303).

### Construction of pseu-dotime trajectory using variable genes

The Monocle2 R package (version 2.10.1) was used to construct single cell pseudo-time trajectories to discover developmental transitions^32–34^. We used highly variable genes identified by “estimateDispersions” function to sort cells in pseudo-time order. “DDRTree” was applied to reduce dimensional space, and the minimum spanning tree on cells was plotted by the visualization functions “plot_complex_cell_trajectory”. BEAM tests were performed on the first branch points of the cell lineage using all default parameters. “plot_genes_branched_pseudotime” function was performed to plot a couple of genes for each lineage.

### Branch expression analysis modeling

Monocle provides the branch expression analysis modeling (BEAM) to calculate the gene expression change as cells pass from early developmental stage through the branch and DEGs between different branches. We used “BEAM” function with default parameter settings and genes with q value less than 0.01 were considered significant.

### Regulatory network construction

We downloaded homo sapiens TF list from AnimalTFDB 3.0^35^ as a TF reference and extracted TFs in marker genes list of each cluster to construct the regulatory network using GENIE3 R package^36^. The network was plotted using Cytoscape.

### Construction of cellular communication network

The ligand-receptor interaction relationships were downloaded from the database, IUPHAR/BPS Guide to PHARMACOLOGY^37^, and the Database of Ligand-Receptor Partners (DLRP)^38,39^. 0.5 was used as a threshold for the average expression level of normalized expression level. Ligands and receptors whose expression above this threshold were considered as expressed in the corresponding cluster. Cytoscape was used to visualize the interactions.

## Supporting information

Table S1: Marker used in cell type assignment

Table S2: DEGs

Table S3: regulatory network in cone PR (age comparison)

Table S4: retinal diseases related genes

## Acknowledgements

We thank the support of a GRF grant from Hong Kong government Project No. [CityU 11256116], a SRG grant from City University of Hong Kong Project No.[7005058], Natural Science Foundation of Guangdong Province (NO.2015A030313472), NSFC grants 81770925 and 81790641 and the Non-profit Central Research Institute Fund of Chinese Academy of Medical Sciences 2018PT32019.

## Author contributions

XH-S, JH-W, SH-Z, F-C, Y-G and DS-C designed and initiated the project; FY-H, JJ-X, P-X, FJ-G, DD-W, DW-Z coordinated and collected the human donor retinas; XH-S, JH-W, SH-Z, and FY-H conducted the experiments of retina sampling; ZX-X, SY-W, LC-L, CC-C and J-X constructed sequencing libraries; JX-Z, SY-W and XN-D were responsible for data analysis; JC-Z, PW-D and W-L contributed to data visualization; JX-Z, DS-C, F-C, JH-W, SJ-J, JG-Z, JK-L, J-F wrote the manuscript and/or provided suggestions. All authors have revised and proofread the manuscript.

## Data and material availability

The data that support the findings of this study have been deposited in the CNSA (https://db.cngb.org/cnsa/) of CNGBdb with accession number CNP0000436.

---

**Figure S1.**
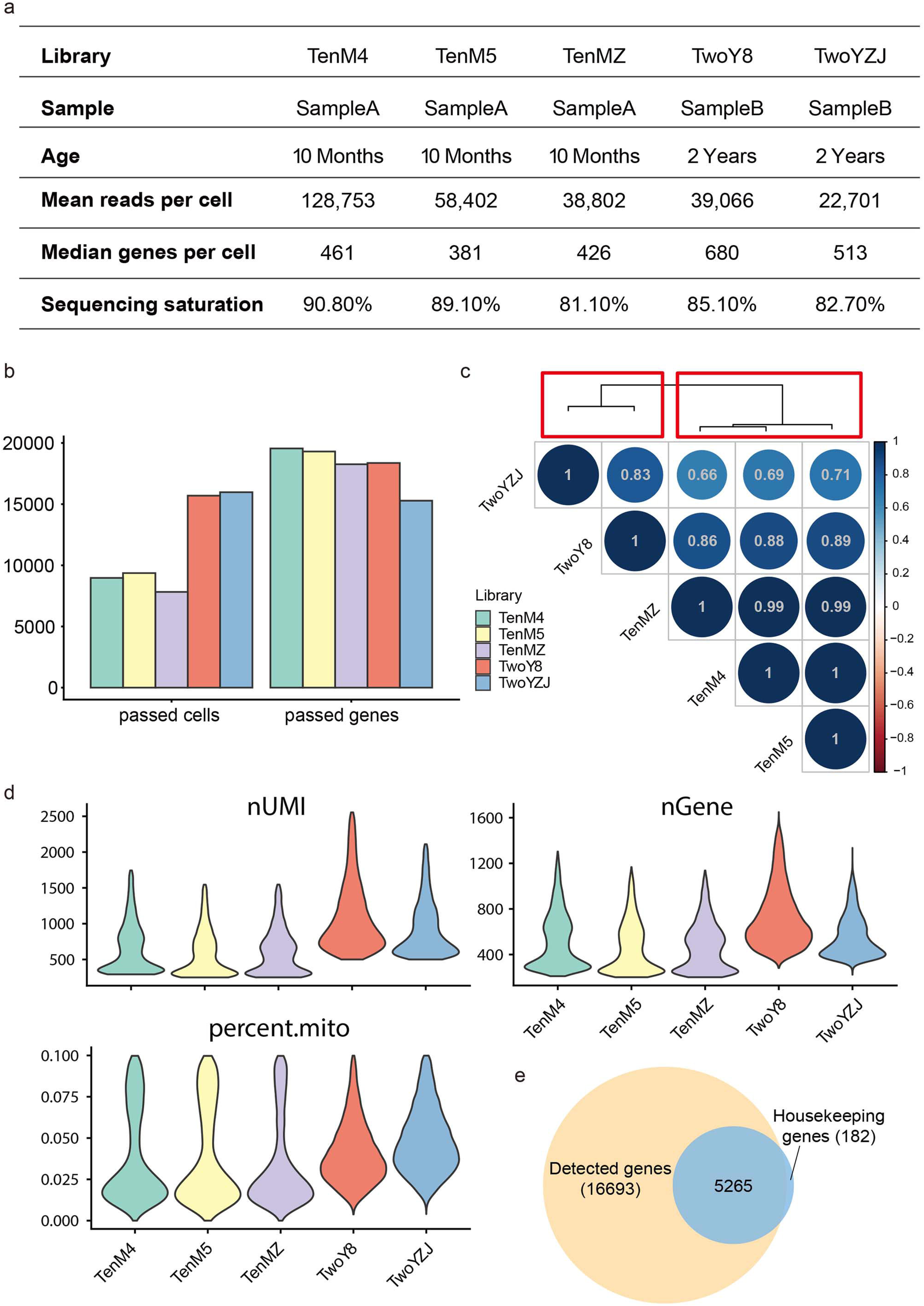
(A) Human infant retinal sampling and sequencing information. (B) Barplot showing the basic library information including the numbers of cells and genes that passed our QC protocol (Methods). (C) Pearson correlation analysis of transcriptomic profiles from all sequencing libraries. Top panel shows the result of hierarchical clustering. TwoY* denote libraries constructed using samples from 2-year old infant while TenM* denote libraries constructed using samples from 10-month old infant. (D) Violin plot showing basic information of each sequencing libraries. (E) Venn plot showing the intersect of housekeeping genes and detected genes in our dataset.

**Figure S2.**
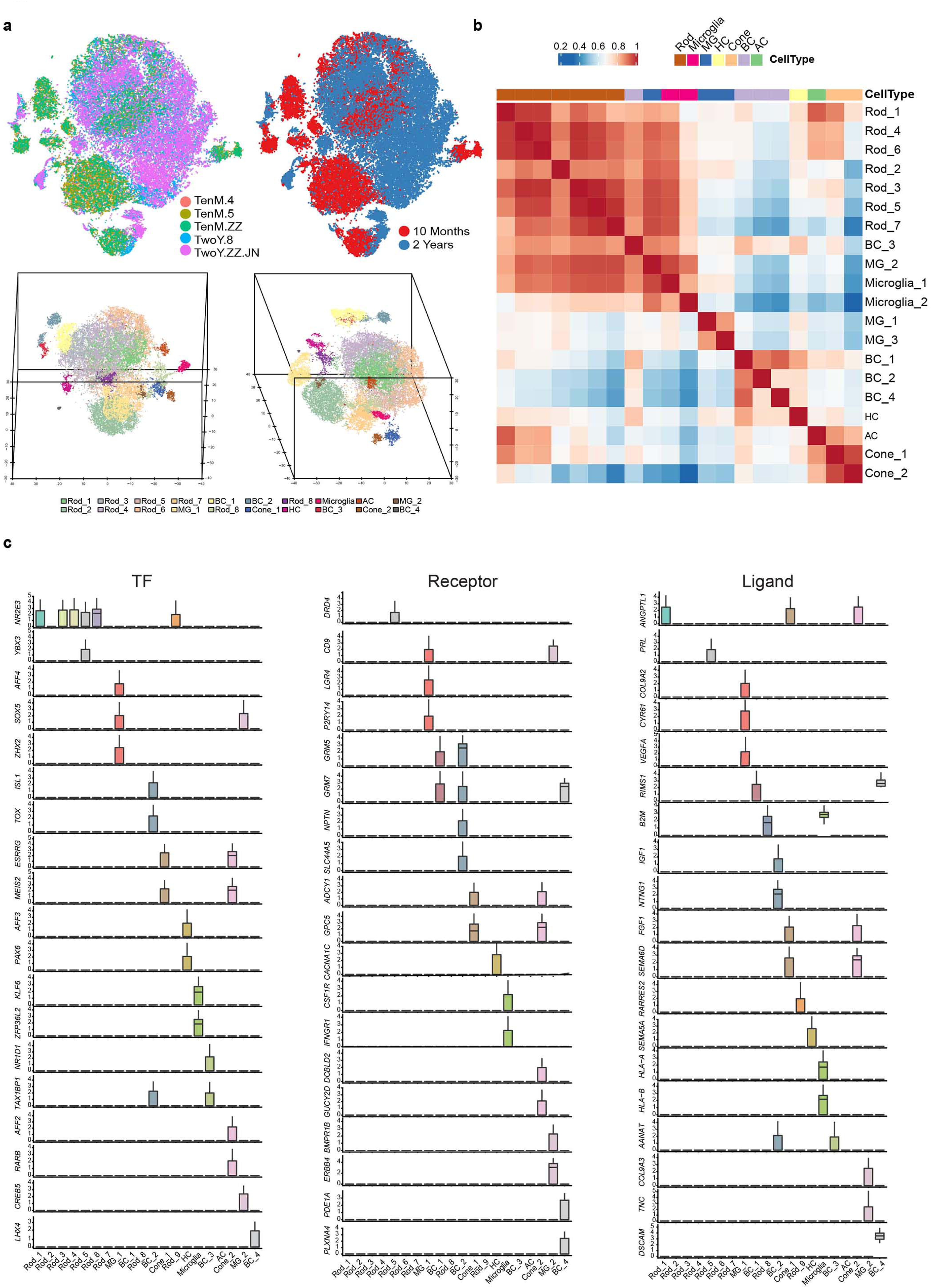
(A) T-SNE maps of the whole dataset. Cells are colored according to their batches (upper left), sample source (upper right). Bottom panel shows 3D t-SNE plots of dimensions 1,2,3 from two different angles. (B) Pairwise correlation coefficients among clusters in Figure 1b. Red indicates high correlation while blue indicates the opposite. (C) Boxplots of differentially expressed TFs, receptors and ligands in each cluster in Figure 1b.

**Figure S3.**
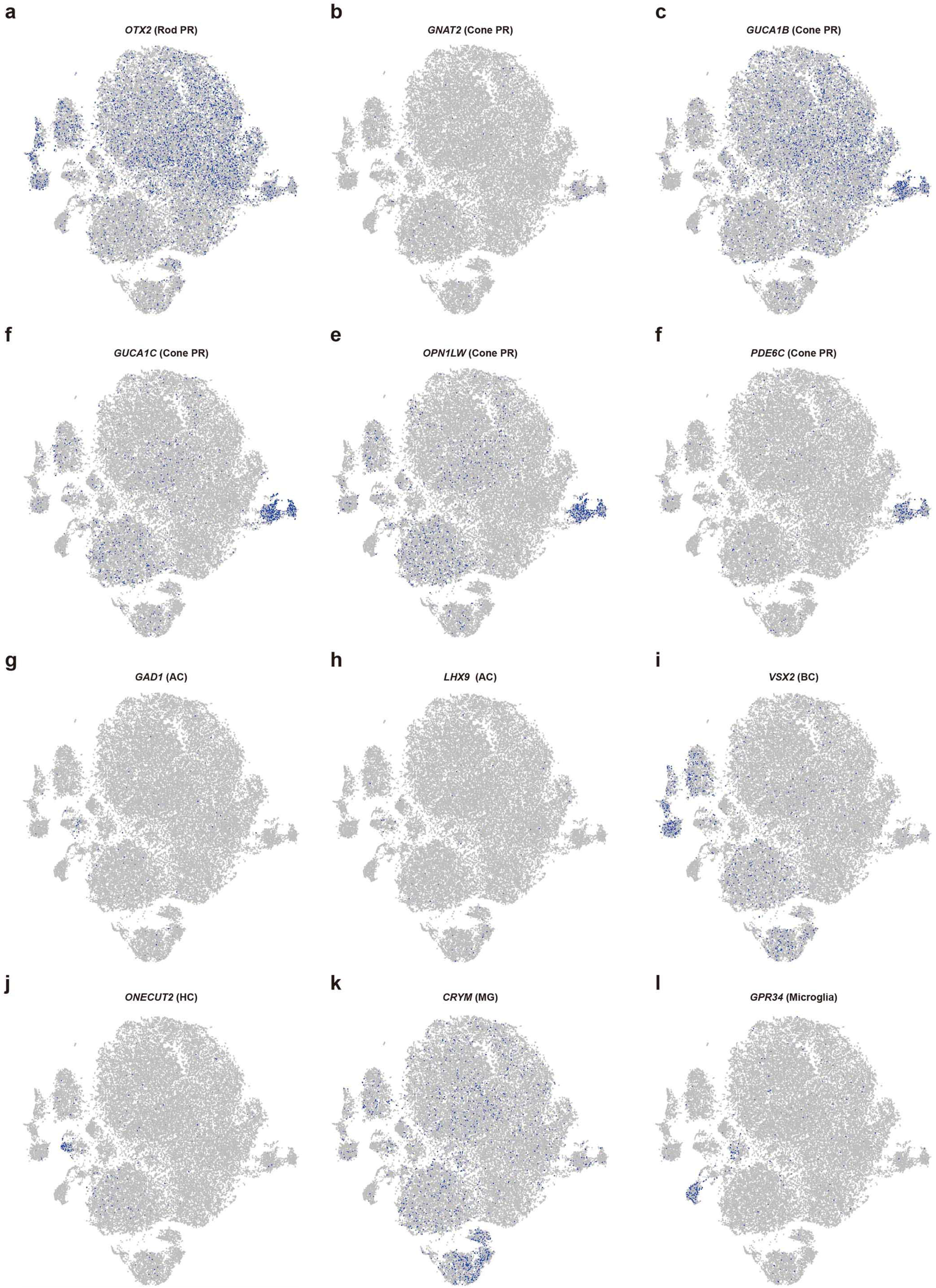
(A-L) T-SNE plots showing the expression pattern of canonical markers for each cell type identified in Figure 1b. Gene expression levels are indicated by shades of blue.

**Figure S4.**
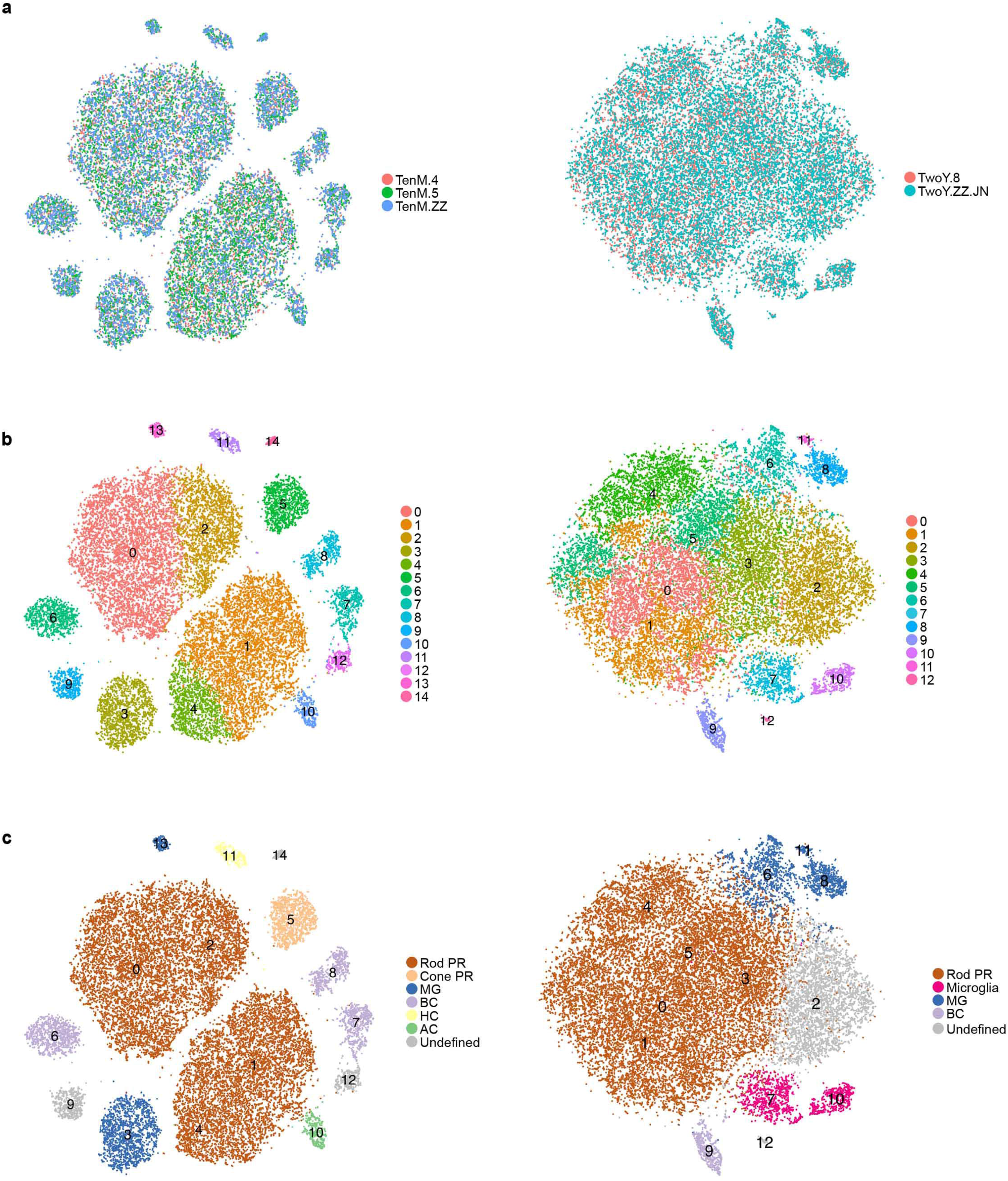
(A) T-SNE plots of individual clustering results of cells from two samples (left: 10 months; right: 2 years). Cells are colored by batches. (B) T-SNE plots of individual clustering results of cells from two samples (left: 10 months; right: 2 years). Cells are colored by clusters. (C) T-SNE plots of individual clustering results of cells from two samples (left: 10 months; right: 2 years). Cells are colored by cell types.

**Figure S5.**
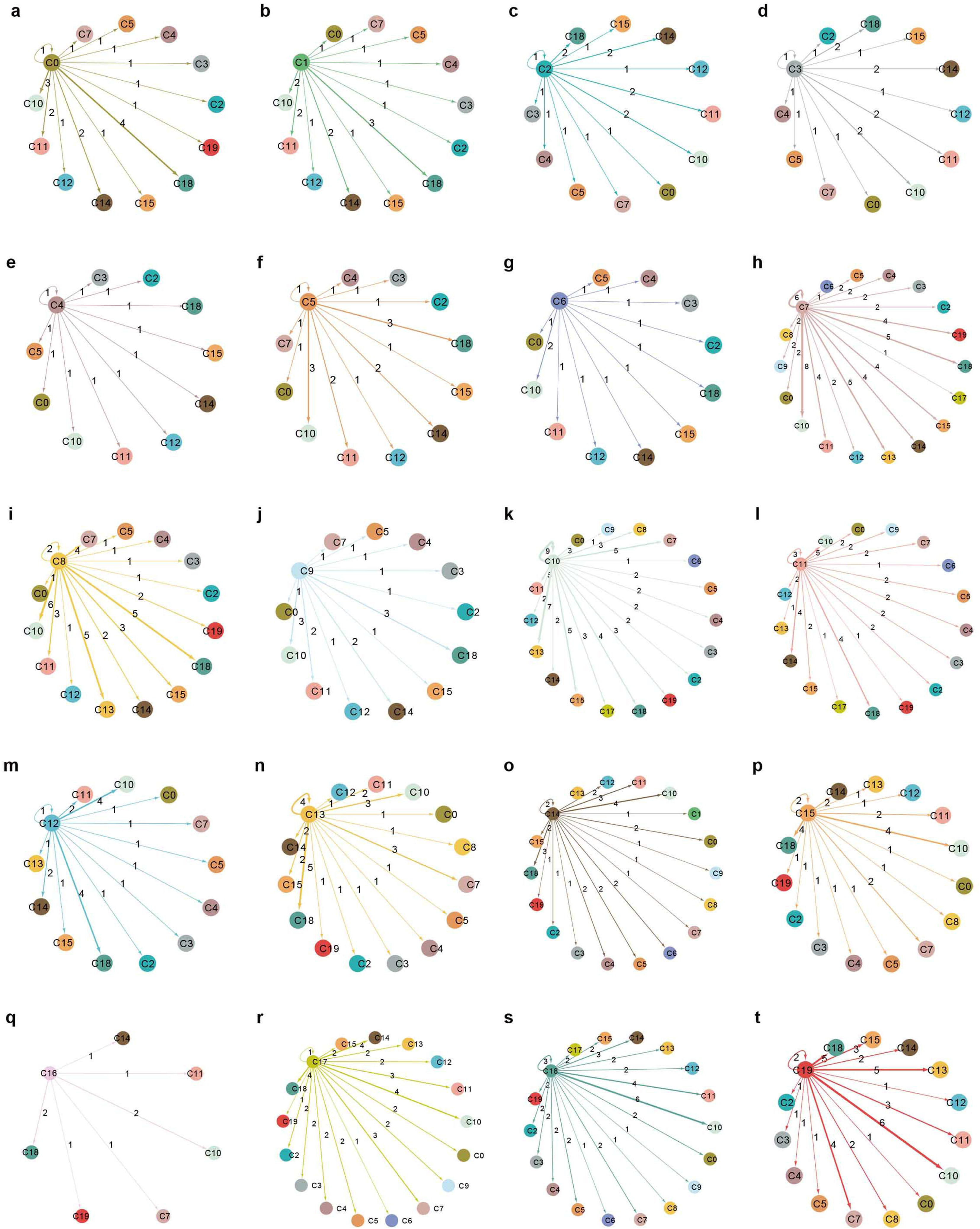
(A-T) Sub-communication network of all clusters in Figure 1b.

**Figure S6.**
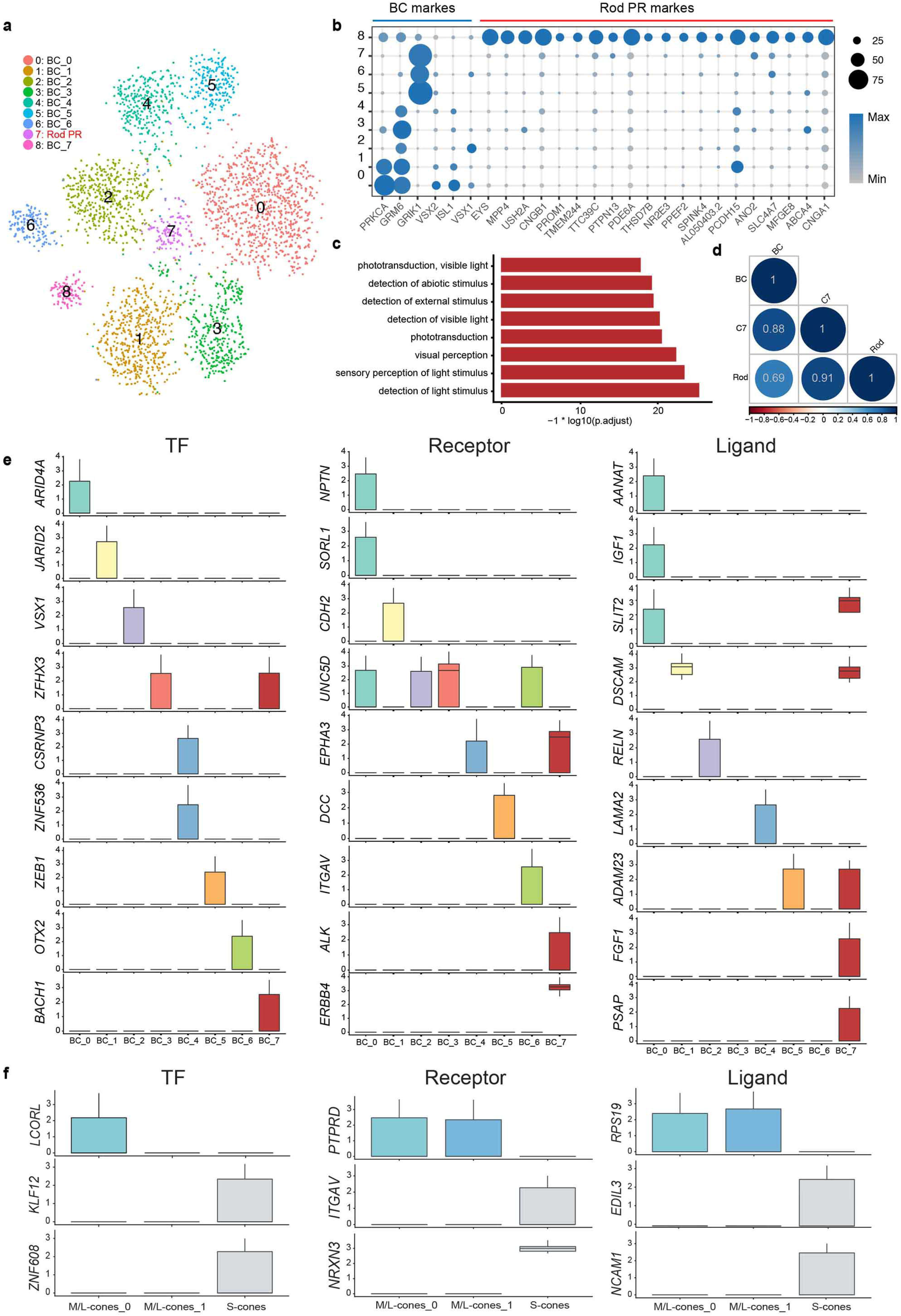
(A) T-SNE plot showing the reclustering result of BC dataset (BC_1, BC_2, BC_3 and BC_4 in Figure 1b). Cells are colored by cluster. C7 were found to be rod photoreceptors. (B) Expression pattern of bipolar cell and rod photoreceptors markers. Dot size represent the percentage of cells express corresponding gene and color represent the expression level. (C) Barplot showing the GO terms enriched in C7 differentially expressed genes. (D) Transcriptomic correlation analysis of C7 between bipolar cells and rod photoreceptors. (E) Boxplots of differentially expressed TFs, receptors and ligands in each cluster in Figure 3g. (F) Boxplots of differentially expressed TFs, receptors and ligands in each cluster in Figure 3a.

**Figure S7.**
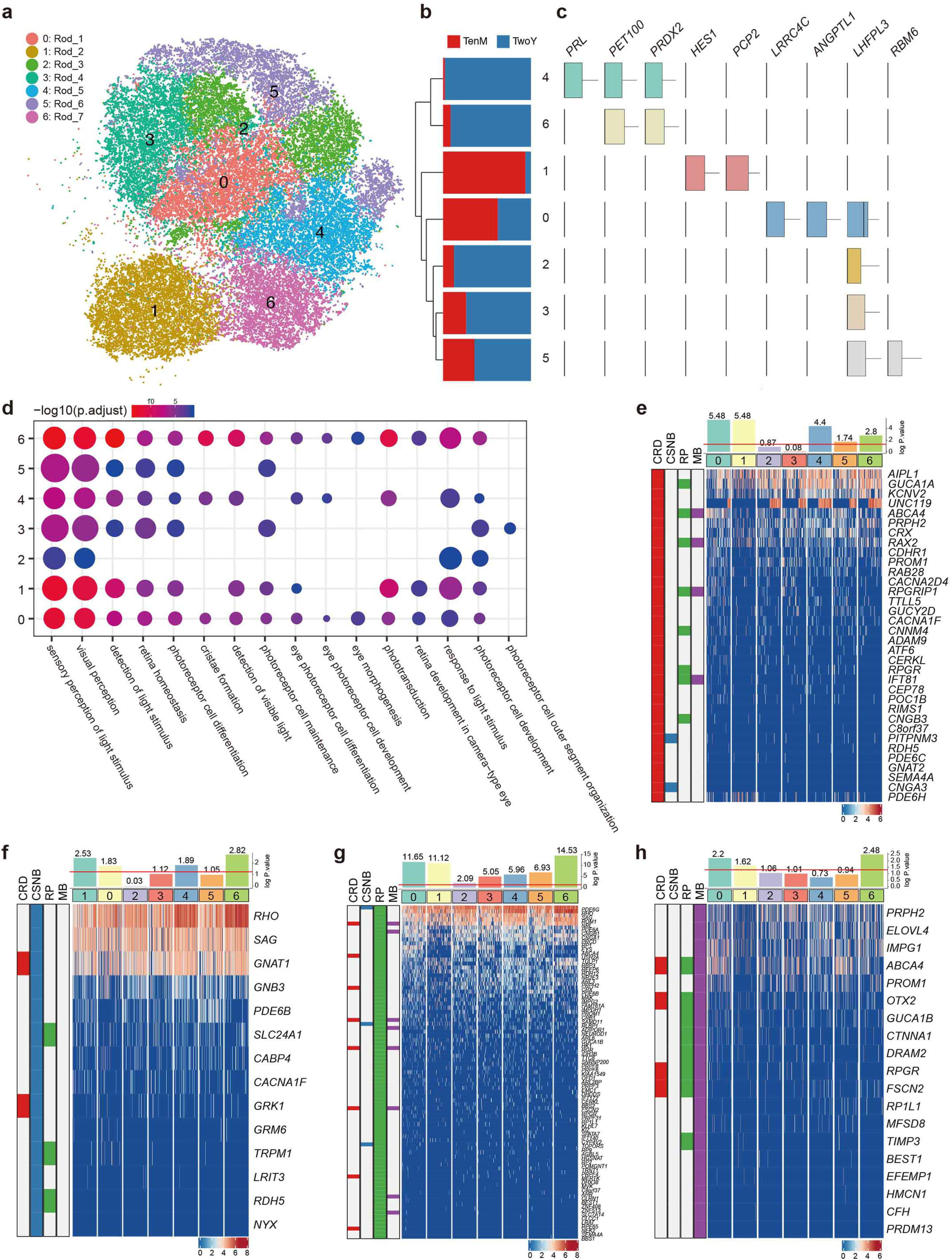
(A) T-SNE plot showing the rod dataset (C0-C6) extracted from Figure 1b. Cells are colored by cluster. (B) Unsupervised hierarchical clustering of the 7 clusters based on the average gene expression of cells in each cluster. The proportions of different cell sources are shown. (C) Boxplots showing the expression patterns of selected differentially expressed genes. (D) Selected GO terms enriched in each cellular cluster. Dot size denotes the gene ratio and color denotes the adjusted P value. (E-H) Rod subpopulation enrichment of CRD (Cone-Rod Dystrophy) (E), RP (Retinitis Pigmentosa) (F), CSNB (Congenital Stationary Night Blindness) (G) and MD (Macular Degeneration) (H) risk genes respectively. Heatmap: Expression of corresponding disease risk genes in rod subpopulations. cells are ordered by cluster. Bar graphs: numbers indicate log2 P value, the red line indicates significance.

**Figure S8.**
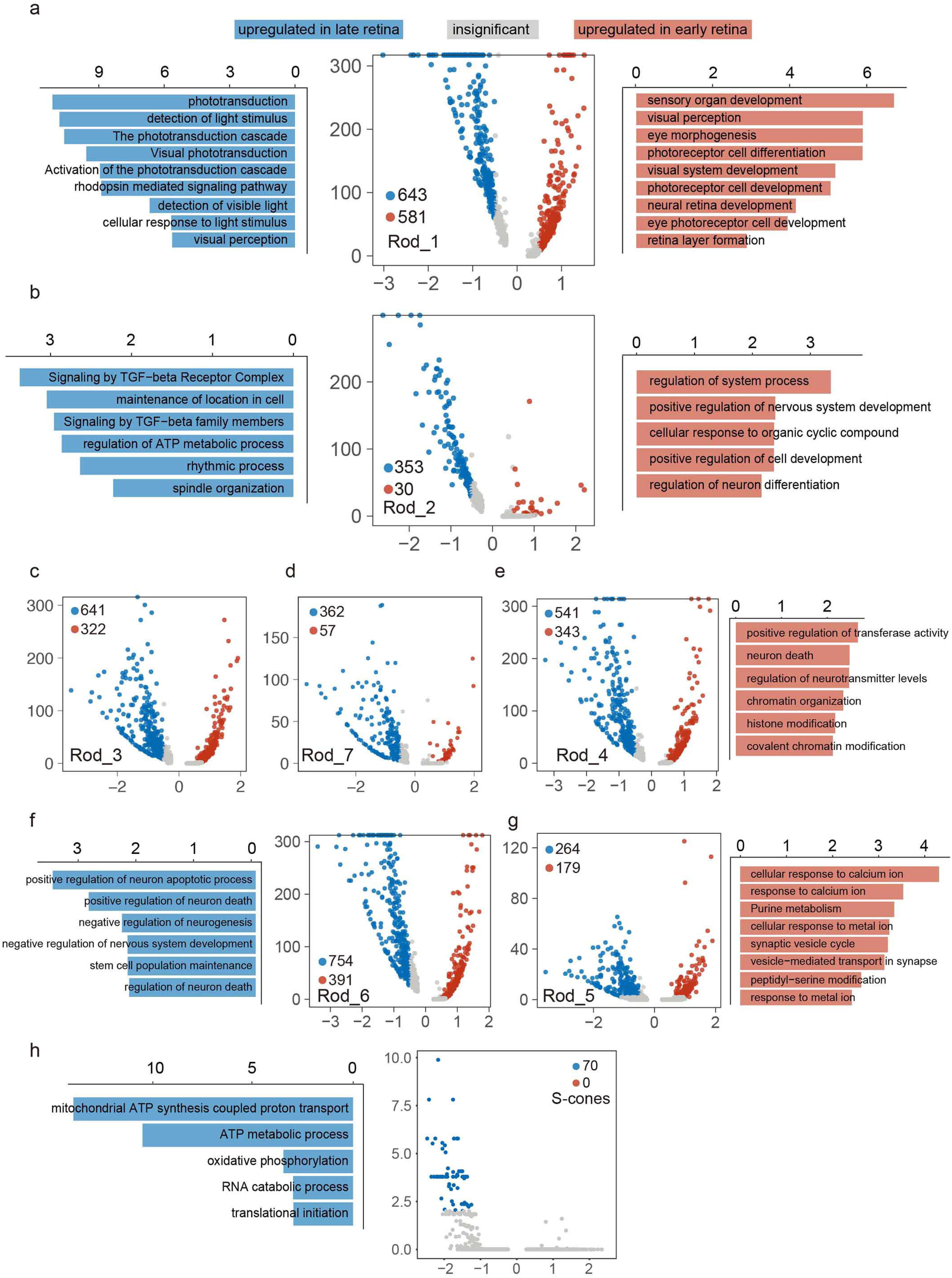
(A-G) Volcano plot: The differentially expressed genes in Rod_1 (A), Rod_2 (B), Rod_3 (C), Rod_4 (E), Rod_5 (G), Rod_6 (F), Rod_7 (D) between early and late retina. Genes are plotted as log2 fold change versus the −log10 of the adjusted p-value. Genes in red denotes genes upregulated in early retina while blue denotes upregulated in late retina. Insignificant genes are colored in grey. Barplot: functional enrichment of genes that significantly upregulated in early (colored in red) or late (colored in blue) retina.

**Figure S9.**
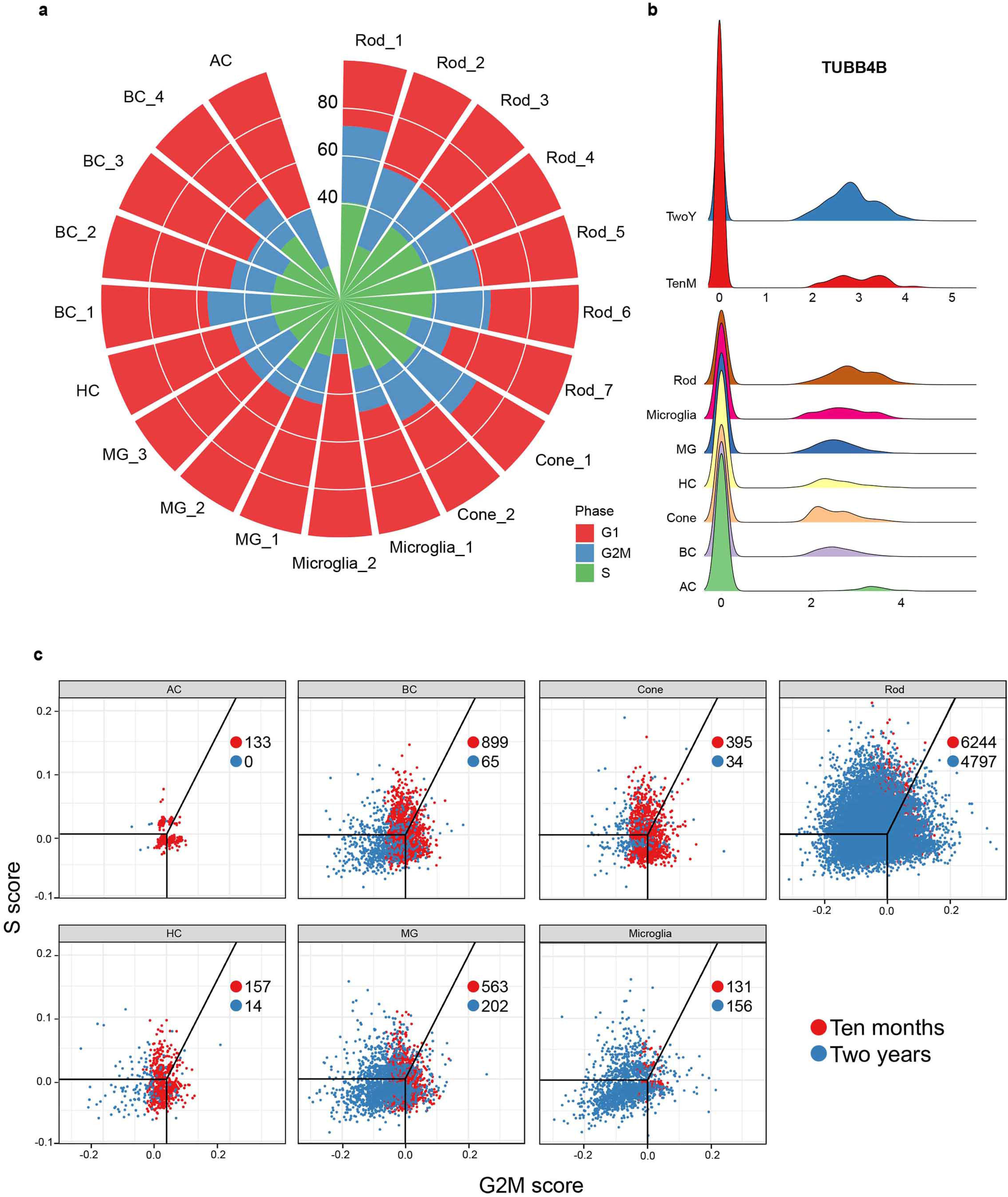
(A) Circular barplot of the percentage of cells in different cell cycles in each cluster in Figure 1b. (B) Ridge plot showing the expression of selected cell cycle related genes in each cell type and sample source. (C) Cell cycle plot for each cell type identified in Figure 1b. Each dot represents a single cell, and its color represents the sample source. The positions of dots illustrate the cell cycle phases of single cells. The inserted numbers represent the numbers of cells in G2M stage in each cell type from different sample sources.

**Figure S10.**
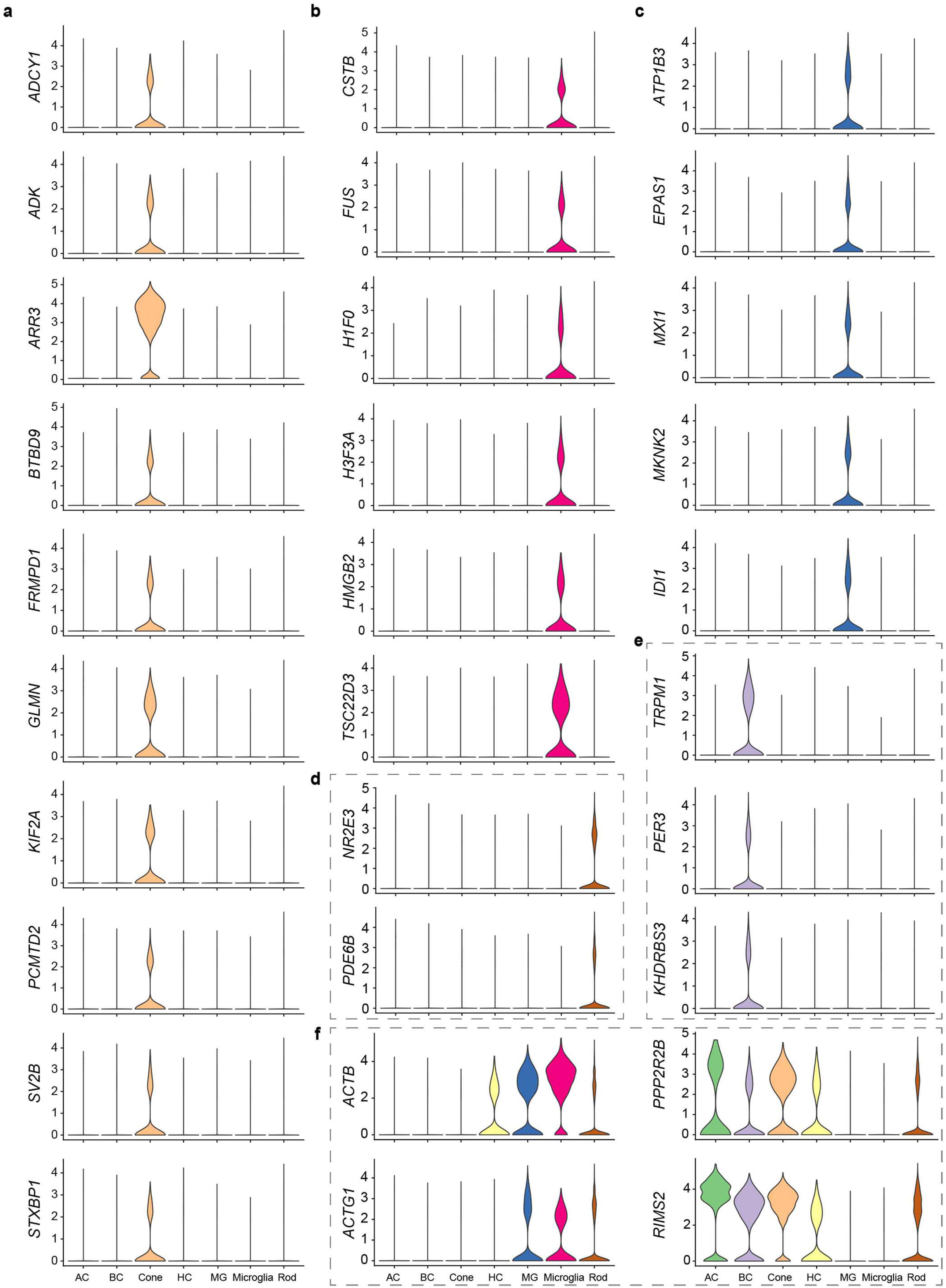
(A-F) Violin plots showing different expression patterns of rhythm related genes in different cell types. Each dash box indicates an expression pattern.

## References

1. Masland, R. H. The Neuronal Organization of the Retina. Neuron 76, 266–280 (2012).

2. Sarthy, P. V. & Lam, D. M. K. Biochemical studies of isolated glial (muller) cells from the turtle retina. J. Cell Biol. 78, 675–684 (1978).

3. Pertile, G., Broccoli, V., Rama, P., Giannelli, S. G. & Demontis, G. C. Adult Human Müller Glia Cells Are a Highly Efficient Source of Rod Photoreceptors. Stem Cells 29, 344–356 (2010).

4. Lukowski, S. et al. Generation of human neural retina transcriptome atlas by single cell RNA sequencing. bioRxiv 425223 (2018). doi:10.1101/425223

5. Rheaume, B. A. et al. Single cell transcriptome profiling of retinal ganglion cells identifies cellular subtypes. Nat. Commun. 9, (2018).

6. Clark, B. et al. Comprehensive analysis of retinal development at single cell resolution identifies NFI factors as essential for mitotic exit and specification of late-born cells. bioRxiv 378950 (2018). doi:10.1101/378950

7. Yufeng Lu, Wenyang Yi, Qian Wu, Suijuan Zhong, Zhentao Zuo, Fangqi Zhao, Mei Zhang, Nicole Tsai, Yan Zhuo, Sheng He, Jun Zhang, Xin Duan, Xiaoqun Wang, T. X. Single-cell RNA-seq analysis maps the development of human fetal retina. bioRxiv (2018). doi:http://dx.doi.org/10.1101/423830

8. Mathis, C., Schröter, A., Thallmair, M. & Schwab, M. E. Nogo-A regulates neural precursor migration in the embryonic mouse cortex. Cereb. Cortex 20, 2380–2390 (2010).

9. Mei, L. & Nave, K. A. Neuregulin-ERBB signaling in the nervous system and neuropsychiatric diseases. Neuron 83, 27–49 (2014).

10. Kung, F., Wang, W., Tran, T. S. & Townes-Anderson, E. Sema3A reduces sprouting of adult rod photoreceptors in vitro. Investig. Ophthalmol. Vis. Sci. 58, 4038–4051 (2017).

11. Singhal, S. et al. Human Müller Glia with Stem Cell Characteristics Differentiate into Retinal Ganglion Cell (RGC) Precursors In Vitro and Partially Restore RGC Function In Vivo Following Transplantation. Stem Cells Transl. Med. 1, 188–199 (2012).

12. Becker, S. et al. Transplantation of Photoreceptors Derived From Human Müller Glia Restore Rod Function in the P23H Rat. Stem Cells Transl. Med. 3, 323–333 (2014).

13. Smith, R. S. et al. Phospholipid flippase ATP8A2 is required for normal visual and auditory function and photoreceptor and spiral ganglion cell survival. J. Cell Sci. 127, 1138–1149 (2014).

14. Lee, A. S. Y., Kranzusch, P. J. & Cate, J. H. D. EIF3 targets cell-proliferation messenger RNAs for translational activation or repression. Nature 522, 111–114 (2015).

15. Hofmann, L. & Palczewski, K. Advances in understanding the molecular basis of the first steps in color vision. Progress in Retinal and Eye Research 49, 46–66 (2015).

16. Neitz, J. & Neitz, M. The genetics of normal and defective color vision. Vision Research 51, 633–651 (2011).

17. Curcio, C. A., Sloan, K. R., Kalina, R. E. & Hendrickson, A. E. Human photoreceptor topography. J. Comp. Neurol. 292, 497–523 (1990).

18. Shekhar, K. et al. Comprehensive Classification of Retinal Bipolar Neurons by Single-Cell Transcriptomics. Cell 166, 1308–1323.e30 (2016).

19. Zenisek, D., Henry, D., Studholme, K., Yazulla, S. & Matthews, G. Voltage-dependent sodium channels are expressed in nonspiking retinal bipolar neurons. J. Neurosci. 21, 4543–50 (2001).

20. Fulton, A. B., Hansen, R. M. & Moskowitz, A. Development of Rod Function in Term Born and Former Preterm Subjects. Optom. Vis. Sci. 86, E653–E658 (2009).

21. Eroglu, E. et al. SWI/SNF complex prevents lineage reversion and induces temporal patterning in neural stem cells. Cell 156, 1259–1273 (2014).

22. Leifer, D. et al. MEF2C, a MADS/MEF2-family transcription factor expressed in a laminar distribution in cerebral cortex. Proc. Natl. Acad. Sci. USA 90, 1546–1550 (1993).

23. Tang, W.-X. et al. The nuclear receptor NR2E3 plays a role in human retinal photoreceptor differentiation and degeneration. Proc. Natl. Acad. Sci. 99, 473–478 (2002).

24. Swaroop, A., Kim, D. & Forrest, D. Transcriptional regulation of photoreceptor development and homeostasis in the mammalian retina. Nature Reviews Neuroscience 11, 563–576 (2010).

25. Zhang, X. et al. Therapeutic Potential of Co-enzyme Q10 in Retinal Diseases. Curr. Med. Chem. 24, (2017).

26. Iijima, K. et al. ZNF385B is characteristically expressed in germinal center B cells and involved in B-cell apoptosis. Eur. J. Immunol. 42, 3405–3415 (2012).

27. Thomas, M. G. et al. Abnormal retinal development associated with FRMD7 mutations. Hum. Mol. Genet. 23, 4086–4093 (2014).

28. Edwards, M. M. et al. Lama1 mutations lead to vitreoretinal blood vessel formation, persistence of fetal vasculature, and epiretinal membrane formation in mice. BMC Dev. Biol. 11, (2011).

29. Baguma-Nibasheka, M. & Kablar, B. Abnormal retinal development in the Btrc null mouse. Dev. Dyn. 238, 2680–2687 (2009).

30. Tirosh, I. et al. Dissecting the multicellular ecosystem of metastatic melanoma by single-cell RNA-seq. Science (80-.). 352, 189–196 (2016).

31. Kowalczyk, M. S. et al. Single-cell RNA-seq reveals changes in cell cycle and differentiation programs upon aging of hematopoietic stem cells. Genome Res. 25, 1860–1872 (2015).

32. Qiu, X. et al. Single-cell mRNA quantification and differential analysis with Census. Nat. Methods 14, 309–315 (2017).

33. Qiu, X. et al. Reversed graph embedding resolves complex single-cell trajectories. Nat. Methods 14, 979–982 (2017).

34. Trapnell, C. et al. The dynamics and regulators of cell fate decisions are revealed by pseudotemporal ordering of single cells. Nat. Biotechnol. 32, 381–386 (2014).

35. Hu, H. et al. AnimalTFDB 3.0: a comprehensive resource for annotation and prediction of animal transcription factors. Nucleic Acids Res. 47, D33–D38 (2019).

36. Huynh-Thu, V. A., Irrthum, A., Wehenkel, L. & Geurts, P. Inferring regulatory networks from expression data using tree-based methods. PLoS One 5, (2010).

37. Harding, S. D. et al. The IUPHAR/BPS Guide to PHARMACOLOGY in 2018: Updates and expansion to encompass the new guide to IMMUNOPHARMACOLOGY. Nucleic Acids Res. 46, D1091–D1106 (2018).

38. Pavličev, M. et al. Single-cell transcriptomics of the human placenta: Inferring the cell communication network of the maternal-fetal interface. Genome Res. 27, 349–361 (2017).

39. Salwinski, L. The Database of Interacting Proteins: 2004 update. Nucleic Acids Res. 32, 449D– 451 (2003).

